# Ordering structured populations in multiplayer cooperation games

**DOI:** 10.1101/021550

**Authors:** Jorge Peña, Bin Wu, Arne Traulsen

## Abstract

Spatial structure greatly affects the evolution of cooperation. While in two-player games the condition for cooperation to evolve depends on a single structure coefficient, in multiplayer games the condition might depend on several structure coefficients, making it difficult to compare different population structures. We propose a solution to this issue by introducing two simple ways of ordering population structures: the containment order and the volume order. If population structure 𝒮_1_ is greater than population structure 𝒮_2_ in the containment or the volume order, then 𝒮_1_ can be considered a stronger promoter of cooperation. We provide conditions for establishing the containment order, give general results on the volume order, and illustrate our theory by comparing different models of spatial games and associated update rules. Our results hold for a large class of population structures and can be easily applied to specific cases once the structure coefficients have been calculated or estimated.

## Introduction

The evolution of cooperation is a fascinating topic that has been studied from different perspectives and theoretical approaches (2, 14, 21, 36). An issue that has led to considerable interest is the extent to which spatial structure allows cooperation to thrive (1, 7, 10, 19, 20, 22, 27, 29-31, 38-40, 46, 48, 51, 53, 55-60). Spatial structure can both enhance cooperation by inducing clustering or assortment (whereby cooperators tend to interact more often with other cooperators (11, 13, 22)) and oppose cooperation by inducing increased local competition (whereby cooperators tend to compete more often with other cooperators (47)). For two-player games or multiplayer games with similar strategies, the balance between these two opposing effects is captured by the “scaled relatedness coefficient” of inclusive fitness theory (31, 46, 59, 60) or the “structure coefficient” of evolutionary game theory (1, 38, 53). These coefficients are functions of demographic parameters, and take into account the degree of assortment, the effects of density dependence, and the strength of local competition resulting from spatial interactions (20, 31, 53). Two different models of spatial structure and associated evolutionary dynamics can be unambiguously compared by ranking their relatedness or structure coefficients: the greater the coefficient, the less stringent the conditions for cooperation to evolve. Hence, different models of population structure can be ordered by their potential to promote the evolution of cooperation in a straightforward way.

Despite the theoretical importance of models leading to a single relatedness or structure coefficient, many examples of social evolution ranging from microbial cooperation (18, 32, 63) to collective action in humans (23, 35, 43) involve games between more than two players with distinct strategies (17, 44). In these cases, the effects of spatial structure cannot be captured by a single coefficient, as higher degrees of association (e.g., “triplet relatedness” (39, 42)) are required to fully describe the condition under which cooperation is favored (49, 59, 62). The need to account for several structure coefficients has so far precluded a simple way of comparing population structures independently of the particular game used to model cooperation.

Here, we propose a framework to order population structures by their potential to promote cooperation that is also valid in the case of games between multiple players with distinct strategies. Our framework allows the comparison of two population structures without referring to any concrete game. We will distinguish two cases, depending on the inclusion relation between the sets of games for which cooperation is promoted under each population structure. (i) The set of games for which the second population structure promotes cooperation is fully contained in the set of games for which the first population structure promotes cooperation (Fig. 1A). In this case, we say that the first population structure is greater than the second in the containment order, and hence a stronger promoter of cooperation. (ii) The set of games for which one population structure promotes cooperation is not fully contained in the set of games for which the other population structure promotes cooperation (Fig. 1B). In this case, we say that the population structure promoting cooperation for a larger volume of games is greater in the volume order, and hence a stronger promoter of cooperation.

**Figure 1.**
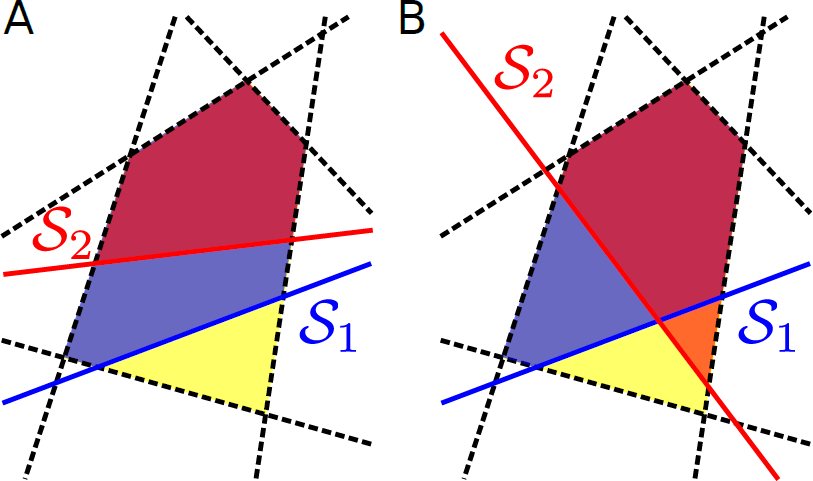
Containment and volume orders of cooperation. The set of *d*-player cooperation games is defined by a set of linear inequalities (*dashed lines*) defining a polytope in a 2*d*-dimensional space. A given population structure (e.g., *S*_1_ or *S*_2_) is characterized by a selection condition defining a further linear inequality (*solid lines*). Here, we show a pictorial representation of the projection of such multidimensional objects to the plane, where polytopes are polygons. (A) The set of games for which cooperation is favored under *S*_2_ is contained in the set of the games for which cooperation is favored under *S*_1_. Hence, we say that Si is greater than *S*_2_ in the containment order (and write 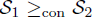). (B) *S*_1_ and *S*_2_ cannot be ordered in the containment order as there are both games for which *S*_1_ favors cooperation but not *S*_2_ (*purple polygon*), and games for which *S*_2_ favors cooperation but not *S*_1_ (*orange polygon*). In both panels, *S*_1_ favors cooperation for more games than *S*_2_ does. Hence, we say that *S*_1_ is greater than *S*_2_ in the volume order (and write 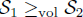).

So far, the structure coefficients for general multiplayer games have been calculated only for few population structures, as such calculations often represent a technical challenge (34). However, once the structure coefficients are known, the containment and volume orders we propose here allow to assess the consequence of population structure on the evolution of cooperation independently of the game at stake. This way, our approach can help to organize myriads of results on the promotion of cooperation in spatially structured populations.

## Methods and Results

### Cooperation games and polytopes

We consider symmetric games between *d* players with two strategies, *A* and *B*. A focal player舗s payoff depends on the player舗s own strategy and on the strategies of its *d* – 1 co-players. If *j* co-players play *A*, a focal *A*-player obtains *a_j_*, whereas a focal *B*-player obtains *b_j_*. These interactions are represented by the payoff table:

| Opposing $A$-players | 0 | 1 | ... | $j$ | ... | $d - 1$ |
| --- | --- | --- | --- | --- | --- | --- |
| payoff to $A$ | $a_0$ | $a_1$ | ... | $a_j$ | ... | $a_{d-1}$ |
| payoff to $B$ | $b_0$ | $b_1$ | ... | $b_j$ | ... | $b_{d-1}$ |

It follows that a game is determined by 2*d* real numbers and can thus be considered as a point in a 2*d*-dimensional space.

In which sense can we say that one population structure favors cooperation more than another? To answer this question precisely, we first need to specify what we mean by “cooperation”, as this could refer to different social behaviors, in particular if we move beyond two-player games (25). We are interested in a particular subset of games that we call “cooperation games”. In these games, players decide whether to cooperate (play *A*) or defect (play *B*), and payoffs are such that: (i) players prefer other group members to cooperate irrespective of their own strategy, and (ii) mutual cooperation is favored over mutual defection. In terms of our payoff parameters, these conditions imply

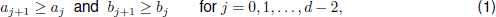

as well as

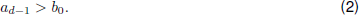

Conditions (1) and (2) are often used to characterize the benefits of cooperation in multiplayer social dilemmas (25), such as the provision of collective goods (46). However, our conditions do not specify individual costs associated to a decision to cooperate, and hence our class of cooperation games includes not only social dilemmas, but also mutualistic games in which individual and group interests are aligned. If we further restrict payoffs to values between zero and one,

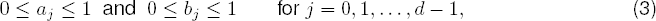

then the set of all cooperation games with *d* players is given by a (convex) polytope (64) in a 2*d*-dimensional space, which we denote by *P*. A polytope is a geometric object with flat sides, the generalization of a polygon (which is a 2-dimensional polytope) to higher dimensional spaces. See Supplementary Material for further details.

We need to specify precisely what we mean by “favoring” cooperation. For our purposes, we say that cooperation is favored if a single cooperator in a population of defectors has a higher probability of eventually reaching fixation than a single defector in a population of cooperators (37). This also means that cooperation is more abundant than defection in a mutation-selection process in the limit of low mutation (16). For weak selection on homogeneous populations of constant size, strategy *A* is favored over *B* if (62)

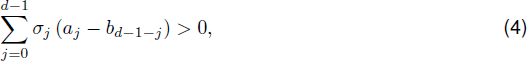

where *σ*_0_…,*σ*_*d*-1_ are the *d* structure coefficients. These are independent of payoffs *a_j_*, and *b_j_*, but dependent on the type of spatial structure (for instance, where the co-players of a given focal individual are located) and update rule used to model the evolutionary dynamics. In Table 1, we provide examples of population structures and their corresponding structure coefficients (see Supplementary Material for a derivation).

**Table 1.**
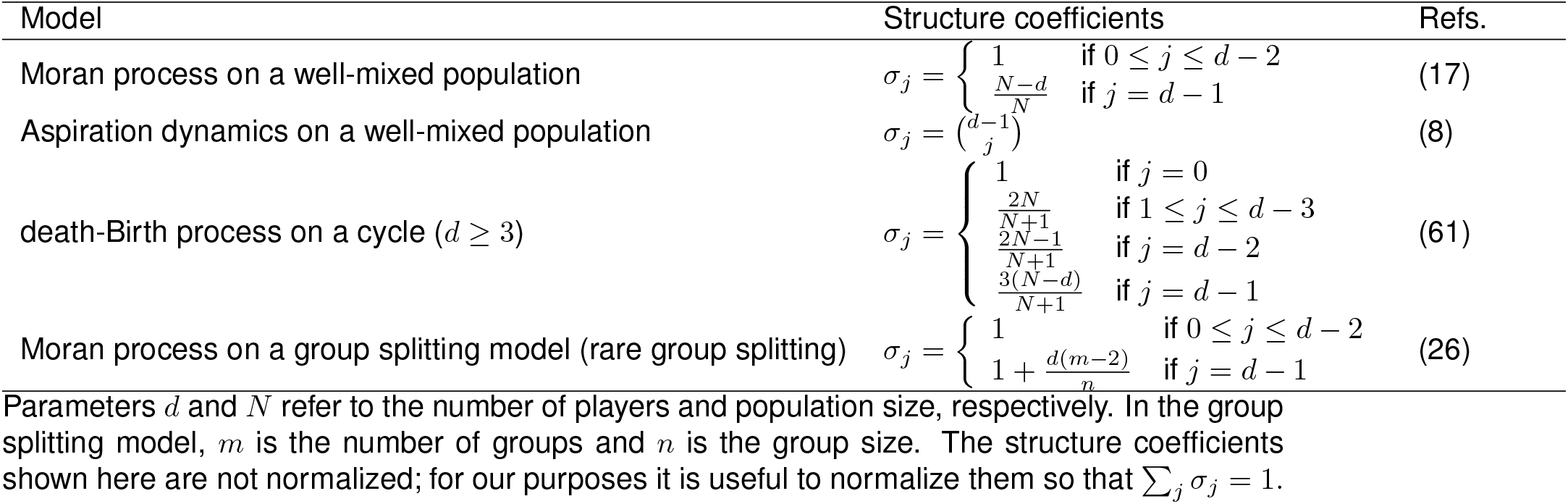
Structure coefficients for some population structures.

The structure coefficients are uniquely determined up to a constant factor. Setting one of them to one thus gives a single nontrivial structure coefficient for two-player games (53). We use the sequence σ to collect the coefficients and note that, if *σ_j_* ≥ 0 for all *j* and *σ_j_* > 0 for at least one *j*, we can impose 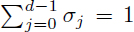 without affecting the selection condition (4). For our purposes, this normalization turns out to be more useful than setting one coefficient to one. In particular, such normalization allows us to understand the (normalized) structure coefficients as describing a probability distribution, and to make a straightforward connection with the concept of assortment as developed for the case of linear public goods games (3, 13). To do so, let us rewrite the selection condition (4) as

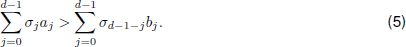

Here, *σ_j_* plays the role of the “effective” probability of interacting with *j* individuals of the own type (and *d* – 1 – *j* of the other type). As given by (5), the selection condition states that *A* is favored over *B* if the expected payoff of an *A*-player is greater than that of a *B*-player when the “interaction environments” (13) are distributed according to *σ*.

A given population structure will favor cooperation only for a subset of cooperation games. More precisely, for a population structure *S_i_* with structure coefficients *σ_i_*, the set of cooperation games for which *S_i_* favors *A* over *B* is given by adding the selection condition (5) to the inequalities defining the polytope of cooperation games, *P*, i.e., (1), (2), and (3). The selection condition (5) defines a hyperplane and thus divides the space of games into two: those for which cooperation is favored and those for which defection is favored. This shows that our problem is equivalent to a geometric problem in 2*d* dimensions. In the following, we denote by *Q_i_* the polytope containing the cooperation games for which cooperation is favored under population structure *S_i_* (see Supplementary Material).

## Containment order

If the set of games *Q*_2_ for which cooperation is favored under population structure *S*_2_ is contained in the set *Q*_1_ for which cooperation is favored under population structure *S*_1_, then we say that *S*_1_ is greater than *S*_2_ in the containment order (12), and we write 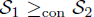. The ordering 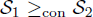 implies that cooperation cannot be favored under *S*_2_ without also being favored under *S*_1_.

Establishing the containment order is equivalent to a “polytope containment problem” (24), consisting on determining whether or not a polytope is contained in another. Polytope containment problems can be solved numerically by linear programming (15). Here, we describe an alternative and simpler approach borrowed from the literature on stochastic orders (50). First, assume that the structure coefficients *σ_j_* are nonnegative and normalized, so that they define a probability distribution over *j* = 0, 1,…, *d* – 1. In this case, the left-hand side of the selection condition (4) can be interpreted as the expected value E[*f*(*J*)], where *f*(*j*) ≡ *f_j_* = *a_j_* – *b*_*d*-1-*j*_, and *J* is the random variable associated to the probability distribution σ. Consider now two population structures *S*_1_ and *S*_2_ with structure coefficients σ_1_ and σ_2_, and associated random variables *J*_1_ and *J*_2_, respectively. A sufficient condition leading to the containment order 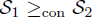 is hence that

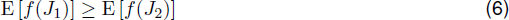

for all cooperation games.

In order to evaluate this condition, we make use of the usual stochastic order (50). A random variable *J*_1_ is said to be greater than *J*_2_ in the stochastic order if and only if 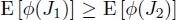 for all increasing functions *ϕ*. This is denoted by 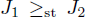. Conveniently, and by (1), the sequence *f_j_* is always increasing in *j*, allowing us to apply this idea directly (see Proposition 1 in Supplementary Material for details). One advantage of expressing the containment order in terms of the stochastic order is that we can transform our original polytope containment problem into the problem of finding conditions under which random variables can be stochastically ordered. Some of these conditions follow from a simple inspection of the sequences of structure coefficients. For instance, a sufficient condition leading to the stochastic order 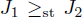 (and hence to the containment order 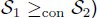) is that σ_1_ – σ_2_ has exactly one sign change from – to + (50). As we show in Examples, this simple condition allows us to order different existing models of population structure in a straightforward way.

For the linear public goods game (i.e., a game with payoffs *a_j_* = β(*j* + 1) – *c* and *b_j_* = *β_j_* for some *β > γ >* 0 where *β* is the marginal benefit from the public good and γ is the individual cost of contributing), the selection condition (5) can be put in a form reminiscent of Hamilton’s rule with (*e*_A_ – *e*_B_) /(*n* – 1) playing the role of a measure of assortment (or relatedness), where 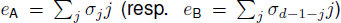 is the mean number of cooperators among the *d* – 1 interaction partners of a cooperator (resp. defector) (3). For more general cooperation games, the selection condition depends not only on the mean but also on higher moments of the probability distribution given by σ. The stochastic order we have used for establishing the containment order is a way of measuring the association between strategies in this general case. Hence, it can be said that population structures greater in the containment order are those characterized by greater “effective assortment” and thus more conducive to the evolution of cooperation. In the extreme case where σ_*d*–1_ = 1 (and *σ_j_* = 0 for *j* ¹ *d* – 1), we have the case of a completely segregated population where *A*s only interact with *A*s and *B*s only interact with *B*s. In this case, the selection condition reduces to (2), and cooperation is always favored by definition.

It can happen that neither *Q*_1_ is entirely contained in *Q*_2_ nor *Q*_2_ is entirely contained in *Q*_1_. In these cases, *S*_1_ and *S*_2_ are incomparable in the containment order (i.e., neither 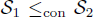 nor 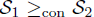 hold) and we write 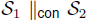. We show in Proposition 2 in Supplementary Material that a sufficient condition leading to such incomparability is that the sequences σ_0_ and σ_2_ cross twice (Fig. 2). In this case, there exist both a subset of cooperation games favored under *S*_1_ but not under *S*_2_ and a subset of cooperation games favored under *S*_2_ but not under *S*_1_.

**Figure 2.**
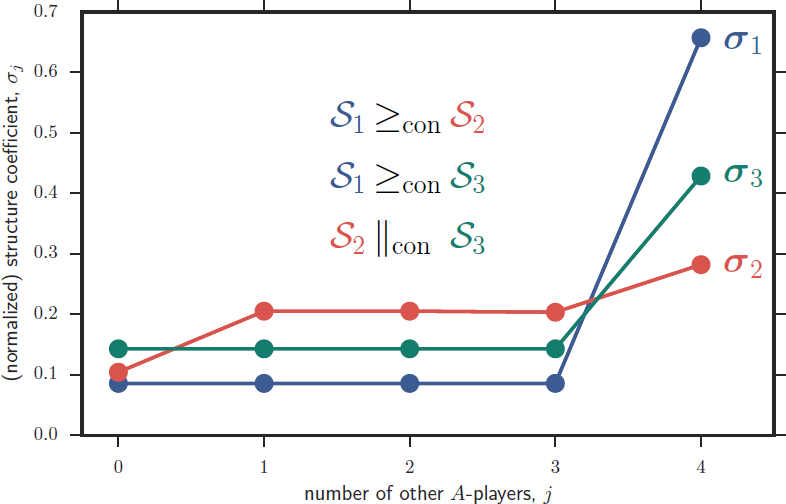
Comparability in the containment order. The structure coefficients σ_1_ and σ_2_ cross exactly once, implying that *S*_1_ and *S*_2_ are comparable in the containment order. Moreover, σ_1_ crosses σ_2_ from below; hence «Si is greater than *S*_2_ in the containment order (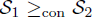). Likewise, *S*_1_ ≥_con_ *S*_3_ Contrastingly, the structure coefficients σ_2_ and σ_3_ cross exactly twice, implying that *S*_2_ and *S*_3_ are incomparable in the containment order (*S*_2_ǁ_con_ *S*_3_, i.e., neither *S*_2_ ≤_con_ *S*_3_ nor *S*_2_ ≥_con_ *S*_3_. For such cases, the volume order provides an alternative way to order these structures. Here, *S*_1_ is a group splitting model with *m* = 10 groups of maximum size *n* = 6 and rare probability of splitting (*q* ≪ 1), *S*_2_ is a cycle of size *N* = 60, and *S*_3_ is a group splitting model with *m* = 6, *n* = 10, and *q* ≪ l.

For the commonly discussed case of two-player games in structured populations (53), the sequence σ consists of two elements: σ_0_ (usually set to one) and σ_0_ (usually denoted by σ and referred to as “the” structure coefficient). Since two sequences of two elements can only cross each other at most once, it follows that any two population structures can be ordered in the containment order if *d* = 2, i.e., the containment order is a total order for two-player games. Moreover, the containment order is given by the comparison of the structure coefficients σ, with larger σ leading to greater containment order. Contrastingly, for *d* ≥ 3 two sequences σ can cross twice. In this case, their respective population structures cannot be compared in the containment order: for multiplayer cooperation games and for the space of all possible population structures, the containment order is only a partial order (see Proposition 3 in Supplementary Material).

### Volume order

In order to address the cases for which two population structures are incomparable in the containment order, we introduce the “volume order”. We say that *S*_1_ is greater than *S*_2_ in the volume order, and write 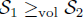, if

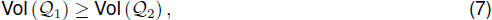

where Vol (*X*) is the volume of polytope *X*. In other words, 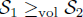 means that for a given *d*, cooperation is favored under *S*_1_ for a greater number of cooperation games than under *S*_2_. If two structures are ordered in the containment order so that 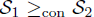, this implies that they are ordered in the volume order so that 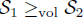, but the converse is not true.

We find that the volume of all *d*-player cooperation games *P* is given by (Proposition 10 in Supplementary Material; Fig. 3):

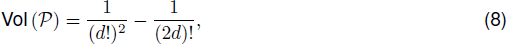

which decreases rapidly with the number of players *d*. For *d* = 2, this volume is equal to 5/24. In this case, the four payoffs *a*_1_, *a*_0_, *b*_1_, and *b*_0_ can be ordered in 4! = 24 possible ways, five of which satisfy inequalities (1) and (2), namely (i) *b*_1_ ³ *a*_1_ ³ *b*_0_ ³ *a*_0_ (prisoner’s dilemma), (ii) *b*_1_ ³ *a*_1_ ³ *a*_0_ ³ *b*_0_ (snowdrift game), (iii) *a*_1_ ³ *b*_1_ ³ *b*_0_ ³ *a*_0_ (stag hunt), (iv) *a*_1_ ³ *b*_1_ ³ *a*_0_ ³ *b*_0_ (harmony game), and (v) *a*_1_ ³ *a*_0_ ³ *b*_1_ ³ *b*_0_ (prisoner’s delight (4)). For large *d*, condition (2) becomes less important and the volume of cooperation games is approximately 1/(*d*!^2^), which is the volume of games satisfying conditions (1) and (3).

**Figure 3.**
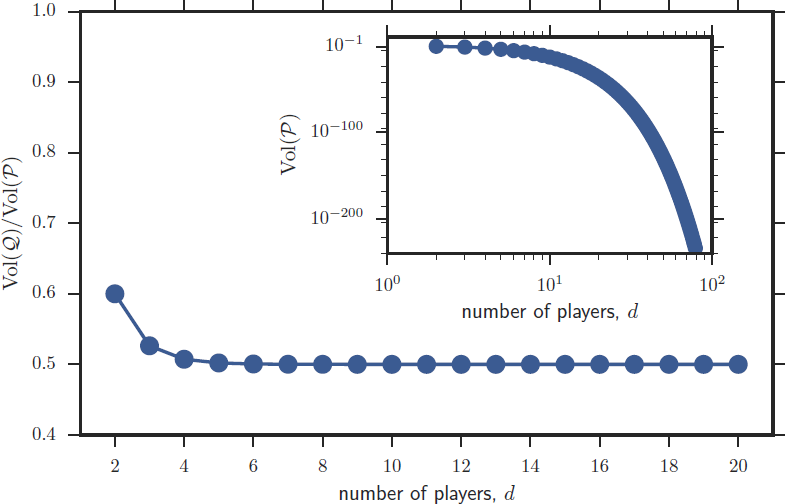
Fraction of cooperation games for which cooperation is favored in population structures with symmetric structure coefficients (*main figure*) and volume of cooperation games (*inset figure*) as functions of the number of players *d.* As *d* increases, the probability that a population structure with symmetric structure coefficients promotes cooperation for a randomly chosen cooperation game quickly approaches 1/2. At the same time, the probability that a randomly chosen game is a cooperation game quickly goes to zero, an effect that seems to be underappreciated in the literature emphasizing the importance of cooperation in evolution.

For some population structures, such as large well-mixed populations updated with a Moran process, the structure coefficients are symmetric, i.e., σ*_j_* = σ_*d*–1–*j*_ for all *j*. For these cases, the fraction of cooperation games for which cooperation is favored becomes

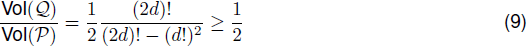

(Proposition 11 in Supplementary Material; Fig. 3). This fraction is equal to 3/5 for *d* = 2, and reduces to 1/2 in the limit of large *d*.

### Examples

Let us now illustrate our approach with particular models of spatial structure and associated update rules. Consider first the baseline scenario of a well-mixed population of size *N* ³ *d* updated with a death-Birth (Moran) process (17, 37). In the death-Birth process, each time step one individual is chosen at random to die and another one is chosen proportional to its payoff to reproduce by making a copy of itself. We find that for any *d* ³ 2, well-mixed populations updated with a death-Birth process are ordered in the containment order with respect to the total population size *N*, such that larger populations are more conducive to multiplayer cooperation (Proposition 4 in Supplementary Material). Our result generalizes previous results for two-player games and multiplayer games with similar strategies according to which smaller population sizes are less conducive to cooperation because of the stronger local competition among cooperators (Ref. (53), Eq. 22; Ref. (60), Eq. B.1). In the limit of large *N* and by Eq. (9), well-mixed populations updated with a death-Birth process favor cooperation for exactly one half of all possible cooperation games.

Consider now the effect of introducing spatial structure while keeping the same update rule. One of the simplest spatial models is the cycle (10). It has been shown that cycles updated with a death-Birth process are better promoters of cooperation than well-mixed populations in the case of two-player games (19, 41), and for several examples of multiplayer social dilemmas (such as linear public goods games, snowdrift games, and stag hunt games) in the limit of large population size (61). Our theory allows us to extend these results to all multiplayer cooperation games and arbitrary population sizes. Indeed, we find that cycles are greater than well-mixed populations in the containment order for any given population size *N* (Proposition 6 in Supplementary Material). This implies that cycles are better promoters of cooperation than well-mixed populations for any cooperation game, any number of players *d*, and any population size *N*.

A second model of spatial structure for which structure coefficients are readily available is the group splitting model of Ref. (58). In this model, a finite population of size *N* is subdivided into *m* groups, which can grow in size and split with probability *q* when reaching the maximum size *n*. In the limit of rare group splitting (*q* ≪ 1), all groups are typically of the maximum size *n* and the structure coefficients can be calculated analytically for general *d*-player games (26). Consider well-mixed and group-splitting populations updated according to a death-Birth process. If the number of groups is greater than two, the group splitting model is greater than any well mixed population in the containment order (Proposition 7 in Supplementary Material). Moreover, in the limit of *m* ≫ *n*, the structure coefficients of the group splitting model become σ_*d*–1_ = 1 and *σ_j_* = 0 for *j* ≠ *d* – 1. In this limit, the group splitting model is greater in the containment order than any other population structure. Hence, it is the population structure that favors cooperation most among all theoretically possible population structures.

The cycle and the group splitting model are better promoters of cooperation than the well-mixed population. But which one promotes cooperation under more cooperation games, the cycle or the group splitting model? Consider cycles of size *N* and group splitting models with rare group splitting (*q* ≪ 1) consisting of *m* groups of maximum size *n*, so that the total maximum population size is equal to *N = mn*. Assuming that the population size *N* is large, the containment order depends on the number of groups *m* of the group splitting model in the following way (Proposition 8 in Supplementary Material). (i) If the number of groups is small (*m* ≤ (*n* + 4*d* – 6)/(2*d* – 3)) the group splitting model is smaller than the cycle in the containment order. (ii) If the number of groups is intermediate (*n* + 4*d* – 6)/(2*d* – 3)) < *m* < *n* + 2) the group splitting model and the cycle are incomparable in the containment order. (iii) If the number of groups is large (*m* ≥ *n* + 2) the group splitting model is greater than the cycle in the containment order. As a particular example, consider a cycle of size *N* = 1000 and a group splitting model with *m* = 10 groups of maximum size *n* = 100 (Fig. 4). In this case, the cycle is greater than the group splitting model in the containment order if *d* ≤ 7, while the two population structures are incomparable in the containment order if *d* ≥ 8. Concerning the volume order, exact computations and numerical simulations suggest that the cycle is greater than the group splitting model for *d* ≤ 12, and smaller than the group splitting model otherwise.

**Figure 4.**
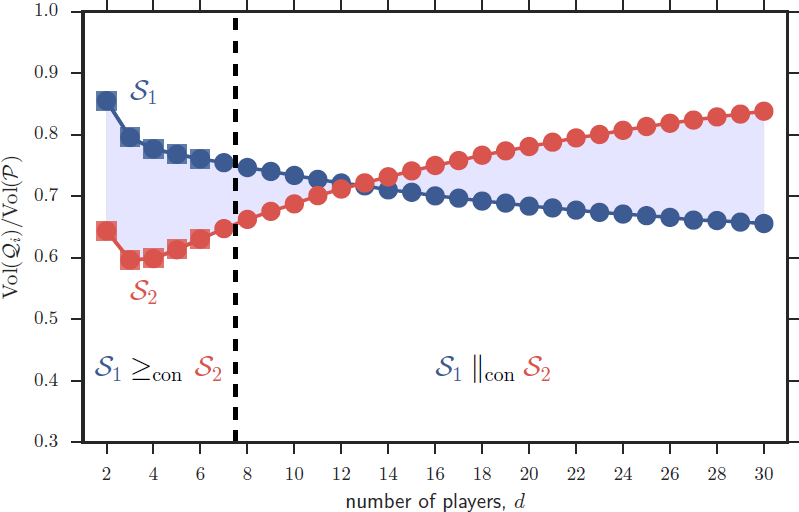
Normalized volumes of cooperation for two different population structures: a cycle of size *N* = 1000 (*S*_1_) and a group splitting model with *m* = 10 groups of maximum size *n*; = 100 (*S*_2_). Volumes are calculated exactly for small values of *d* (squares) and approximately using a Monte Carlo method (*circles*); see Appendix. The cycle is greater than the group splitting model in the volume order for *d* ≤ 12 and smaller in this sense for *d* ≥ 13. We can also show that the cycle is greater than the group splitting model in the containment order (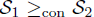) for *d* ≥ 7, but the two population structures are incomparable in the containment order 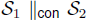 for *d* ≥ 8.

Up until now we have compared different models of spatial structure (the well-mixed population, the cycle, the group splitting model) with a single update rule (the Moran death-Birth process). However, the structure coefficients depend both on spatial structure and on the update rule. For two-player games, different update rules can have important consequences on the evolutionary dynamics, as they lead to different “circles of compensation”, or how far the effects of density dependence extend from a given focal individual (20). What are the effects of different update rules on multiplayer cooperation games? As an example, consider well-mixed populations with two different update rules: the Moran process, where a random individual dies and its neighbors compete for the empty site, and the aspiration dynamics, where an individual is likely to switch its strategy if the current payoff does not meet an aspiration level (8, 52). The two update rules can be ordered in the containment order only if 2^*d*–1^ (*N–d*) ≤ *d*(*N*–1) (Proposition 9 in Supplementary Material). In this case, aspiration dynamics is greater in the containment order than the Moran process, meaning that if cooperation is favored under the Moran process it will also be favored under aspiration dynamics, but not necessarily vice versa. If 2^*d*–1^ (*N – d*) > *d*(*N* – 1) the two structures are incomparable in the containment order. However, for any finite population size *N*, aspiration dynamics is greater in the volume order: overall, cooperation is favored for more games under aspiration dynamics than under the Moran process.

### Discussion

Our approach to compare models of population structure sheds new light on how to study and analyze the evolution of cooperation in spatially structured populations. We have shown how several existing results, obtained under the assumptions of pairwise interactions, similar strategies, or particular classes of multiplayer social dilemmas, generalize to the case of multiplayer cooperation games with distinct strategies that we have considered here. Perhaps more importantly, one can find two population structures such that there is a class of cooperation games for which cooperation is favored under the first but not under the second, and a class of cooperation games for which the opposite holds true (Fig. 1B). Thus, arbitrarily choosing one or a few games from the set of all possible cooperation games to compare the effects of population structure on the evolution of cooperation can be misleading, even when focusing on the comparison of fixation probabilities under weak selection. This is different from the case of either two-player games or multiplayer games with similar strategies, where a ranking of population structures is always possible in terms of a single real value, and where it is sufficient to focus on a single game without loss of generality (31, 53).

We made use of the theory of stochastic orders (50) to provide conditions under which two population structures are comparable or incomparable in the containment order. Within social evolution theory, stochastic orders have been also recently used to tackle the question of whether variability in the group size distribution would lead to less stringent conditions for the evolution of cooperation in multiplayer social dilemmas (45). Our use of stochastic orders in this paper relies on the assumption (fulfilled by all the population structures we used as examples) that the structure coefficients can always be normalized to define a probability distribution. It would be interesting to investigate under which general conditions such assumption is valid. Another open question is whether two population structures incomparable in the containment order could favor cooperation in disjoint subsets of cooperation games. If the structure coefficients define a probability distribution, this will never be the case, as it will always be possible to find a cooperation game for which the selection condition holds for any two population structures. Consider for instance a game for which *a_j_* = α and *b_j_* = 0 for all *j*, with α > 0 (a mutualistic game where the group optimal action *A* is also individually optimal). In this case, and provided that the structure coefficients are nonnegative, the selection condition (4) is always satisfied.

We considered a very broad definition of cooperation and a particular measure of evolutionary success, and investigated subset containment relations and volumes of the resulting polytopes. In this respect our approach is related to a classical study by Mattesi and Jayakar (33), who first defined “analtruism domain” from a set of linear inequalities involving “local fitness functions” and then investigated the problem of finding and measuring the relative volume of the “subset of the altruism domain in which *A* is more fit than *B* on average, that is, altruism can evolve”. We note, however, that our definition of cooperation is different from the definition of altruism adopted by Matessi and Jayakar: the “multi-level interpretation” of altruism, in the sense of Kerr et al. (25). In particular, we only focused on the group benefits, not the individual costs, associated to expressing the cooperative action *A*. Such costs could be introduced by adding further sets of inequalities to the ones we used here, for instance by requiring that *a_j_* ≤ *b_j_* for some or all *j* (25, 46). Since we did not specify any costs, our class of cooperation games contains a relatively large set of mutualistic games for which group beneficial behaviors are also individually beneficial. Our measure of evolutionary success is also different, as we focused on the comparison of fixation probabilities in the limit of weak selection, whereas Matessi and Jayakar focused on the one-step change in frequency. Finally, Matessi and Jayakar limited themselves to “linear fitness functions” (equivalent to linear games in our setup) while we considered more general multiplayer games. The differences between our study and the one by Matessi and Jayakar pinpoint possible future work along these lines. For instance, alternative definitions of cooperation that take in consideration the cost of cooperation (6, 25) and exclude mutualistic games could be explored, possibly together with alternative measures of evolutionary success (54). As long as it is possible to write all conditions as a set of linear inequalities (and hence as polytopes) involving the payoffs of the game, our definitions can be used and adapted to these cases. It would be interesting to see the extent to which comparisons of different population structures based on the containment and volume orders defined here are robust to changes on the way cooperation and evolutionary success are defined and implemented.

### Appendix

#### Computing volumes

There are many exact methods for computing volumes of polytopes, including triangulation methods (5) and signed decomposition methods (28). Computing the exact volume of a polytope is however known to be #P-hard (9), and a simple task only for low dimensions. We calculated exactly the volumes in Fig. 4 for *d* ≤ 6 using the function `volume` of the class `Polyhedron` of the mathematics software Sage (version 6.5). For *d* ≥ 7, we used a Monte Carlo method for approximating the volumes. For each value of *d*, we randomly generated 10^6^ increasing sequences *a_j_*, and *b_j_*, and retained only those which fulfilled (2). We then checked how many of these sequences verified the selection condition (4). The fraction of cooperation games was then approximated by the ratio between these two numbers. Our source code in Python is publicly available on GitHub (https://github.com/jorgeapenas/ordering).

### Supplementary Material

Supplementary Methods: Ordering structured populations in multiplayer cooperation games (PDF).

## Acknowledgments

We would like to thank Laurent Lehmann, Georg Noldeke, and three anonymous referees for their valuable comments on previous versions of this paper.

### Authors’ Contributions

J.P. designed the study and wrote the code; J.P. and B.W. performed research; all authors wrote the manuscript.

### Funding Statement

J.P. acknowledges funding support by the Swiss National Science Foundation (grant PBLAP3-145860).

## Supplementary Material Supplementary Methods: Ordering structured populations in multiplayer cooperation games

### 1 Polytopes

A (convex) polyhedron can be defined as the intersection of finitely many closed halfspaces in ℝ^n^, i.e., as the set of solutions to a system of m linear inequalities

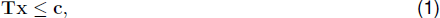

where T is a real *m × n* matrix, and c a real vector of size *m*. A (convex) polytope is a bounded polyhedron. When given by a system of linear inequalities such as (1), a polytope is said to be given in its *H*-representation (18).

We consider symmetric games with two pure strategies (*A* and *B*) between *d* players. A focal player’s payoff depends on its own strategy and on its *d* – 1 co-players’. If *j* co-players play *A*, a focal player obtains *a_j_* if it plays *A* or *b_j_* if it plays *B*. We focus on a subset of games that we call “cooperation games”. In these games *A* represents cooperation; *B*, defection; and payoffs are such that: (i) irrespective of its own strategy, a focal player prefers co-players to cooperate, and (ii) mutual cooperation is favored over mutual defection. In terms of the payoffs of the game, these conditions respectively imply

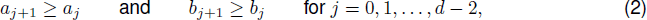

and

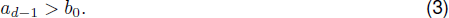

We further restrict payoffs to values between 0 and 1, so that

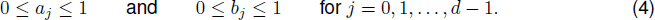

The previous inequalities give the *H*-representation of a polytope that we denote by *P*.

For weak selection, strategy *A* is favored over *B* if (17).

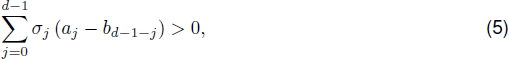

where *σ*_0_,…,*σ_d-1_* are the *d* structure coefficients of the population structure (and associated update rule) under consideration. For a given population structure *S_i_* with vector of structure coefficients *σ_i_*, inequalities (2), (3), and (4) together with the selection condition (5) give the *H*-representation of a polytope that we denote by *Q_i_*.

### 2 Structure coefficients

The structure coefficients σ of a given model of population structure can be calculated from the condition *ρ_A_* > *ρ_B_*, where *ρ_X_* denotes the fixation probability of a single mutant playing *X* (either *A* or *B*) in a population of residents playing the opposite strategy (9). This condition can be written in terms of selection coefficients (dependent on the payoffs of the game and the demographic parameters of the model) and expected coalescence times under neutrality (12). However, the expected coalescence times required for calculating the structure coefficients of general *d*-player games can be difficult to obtain (5, 12). At least for simple population structures, the condition *ρ_A_* > *ρ_B_* can be more easily calculated from first principles, and the structure coefficients extracted from the resulting expressions. This is the approach we followed here.

#### 2.1 Moran process on a well-mixed population

The condition *ρ_A_* > *ρ_B_* is given by (ref. (2), Eq. S19)

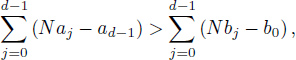

which can be rewritten as

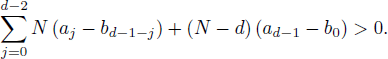

Dividing both sides of the inequality by *N* and comparing with the selection condition (5), we obtain

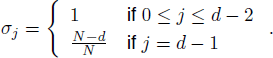

#### 2.2 Aspiration dynamics on a well-mixed population

The condition *ρ_A_* > *ρ_B_* is given by (ref. (1), Eq. 3.3)

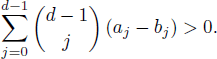

Due to the symmetry of binomial coefficients 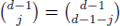, this can be rewritten as

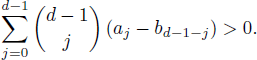

Comparing this last expression with the selection condition (5), we obtain

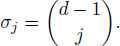

#### 2.3 death-Birth process on a cycle

Let us consider the model of population structure discussed in ref. (14). Each individual is placed on the node of a cycle. Every sequence of *d* players defines the participants in a *d*-player game. Individuals accumulate the payoffs from the *d* games they are involved in, each with *d* players. These payoffs are transformed to “fitness” via an exponential payoff-to-fitness mapping (11). Each time step, a randomly chosen individual is selected to die and its two neighbours compete for the vacant spot with a probability proportional to fitness.

If the population starts with a single mutant, mutants form a single connected cluster in the cycle at any time. The state of the population can hence be captured by the number of *A*-players in this cluster, *i*. Denote by 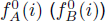 the payoff of an *A*-player (*B*-player) lying immediately at the boundary of a cluster of *B*-players (*A*-players), and by 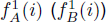 the payoff of an *A*-player (*B*-player) right next to it (Fig. 1). Let 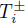 be the probability that the number of *A*-players increases (+) or decreases (-) in one time step, when there are *i A*-players in the population. Hence 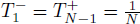

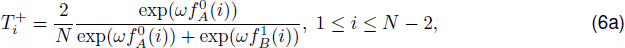

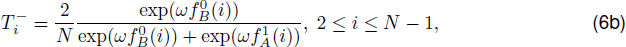

where *ω* is a parameter measuring the strength of selection. Strategy *A* is favored if *ρ_A_* > *ρ_B_*, which is equivalent to (6)

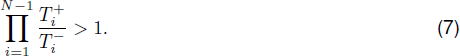

For weak selection, condition (7) is equivalent to

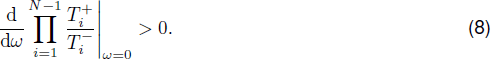

Replacing (6) into (8), we get

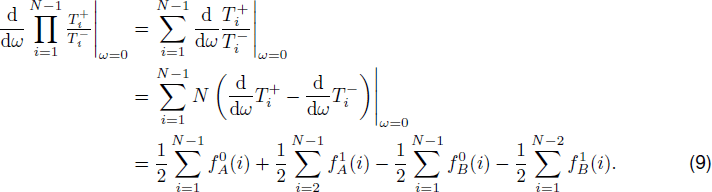

**Figure 1.**
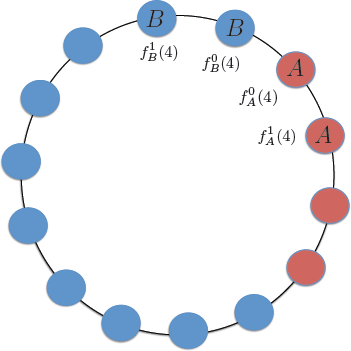
Example of the death-birth process on a cycle. The population is embedded on a cycle of size *N* = 14. There are *i* = 4 *A*-players. 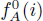 (resp. 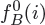 is the payoff of an *B*-player (resp. 5-player) next to the boundary between *A*-players and *B*-players. 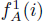 (resp. 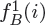) is the payoff of an *A*-player (resp. *B*-player) second-to-next to the boundary between *A*-players and *B*-players.

The above expression is a linear combination of the payoff entries *a_i_* and *b_j_*. Further, by the selection condition (5), the coefficients of *a*_j_ are the same as those of *b_d-1-j_*. Thus it is only necessary to calculate the coefficients of *a*_j_, which only depend on the first two terms of (9). Using the expressions for 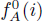 and 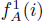 (see ref. (14), Appendix B) we finally obtain:

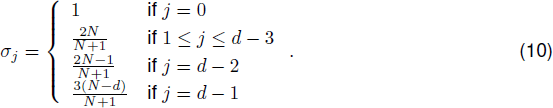

It is noteworthy that the expression

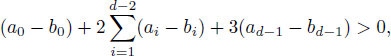

obtained in ref. (14) is a sufficient condition for strategy *A* to be more abundant than strategy *B*. Our result (5) with structure coefficients given by (10) is the necessary and sufficient condition. In addition, we note that our result holds for general payoff-to-fitness mapping *f*, provided that *f’(0)* is non-vanishing (16). Finally, we note that the stricture coefficients given in (10) are valid for *d* ≥ 3. For *d = 2*, they are given by σ_0_ = 1 and σ_1_ = *(3N – 8)/N* (see Eq. 38, Ref. (9)).

#### 2.4 Moran process on a group splitting model

Consider the following multiplayer extension (4) of the group splitting model of ref. (10). The population is subdivided into *m* groups. Each population is allowed to grow to its maximum size *n*, then splits with probability *q*. Within populations, random groups of *d* individuals form and interact in a *d*-player game. When group splitting is rare (*q* ≪ 1) and the mapping between payoffs and fitness is given by an exponential function, the ratio of fixation probabilities is given by (ref. (4), Eq. 15):

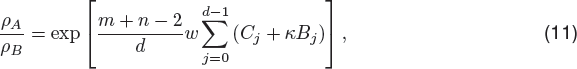

where

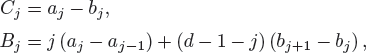

are, respectively, the “direct” and “indirect” gains from switching from strategy *A* to strategy *B* (see ref. (7), Eqs. 6 and 7), and

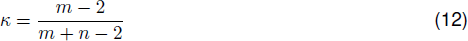

can be interpreted as the “scaled relatedness coefficient” of this model when the migration rate is zero (see ref. (13), Eq. B.4).

From (11), a necessary and sufficient condition for *ρ_A_ > ρ_A_* is that:

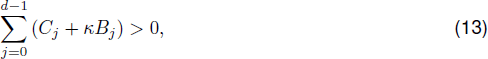

which can be rearranged in terms of payoffs and structure coefficients in the form of the left hand side of (5):

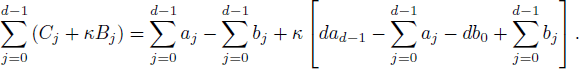

Hence, condition (13) can be written as:

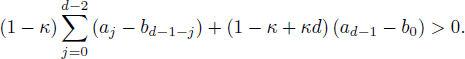

After dividing both sides of the inequality by *1 — κ*, inserting the value of *κ* given in (12), and comparing with (5), we obtain

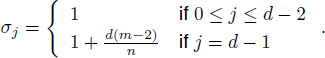

#### 2.5 Normalized structure coefficients

For all population structures discussed in this section and listed in Table 1 of the main text, the structure coefficients are nonnegative. This is also true for many other population structures, at least for *d = 2* (9). In these cases, the structure coefficients can be normalized so that the containment order can be investigated using stochastic orders (8). Henceforth, we refer to the normalized structure coefficients by σ = *(σ_0_,…,σ_d-1_)*. Table 1 lists the normalized structure coefficients for the examples of population structures previously discussed.

**Table 1.** Normalized structure coefficients for multiplayer evolutionary game models.

| Model | Normalized structure coefficients |
| --- | --- |
| Moran process on a well-mixed population | $\sigma_j = \begin{cases} \frac{N}{d(N-1)} & \text{if } 0 \leq j \leq d-2 \\ \frac{N-d}{d(N-1)} & \text{if } j = d-1 \end{cases}$ |
| Aspiration dynamics on a well-mixed population | $\sigma_j = \frac{\binom{d-1}{j}}{2^{d-1}}$ |
| death-Birth process on a cycle ($d \geq 3$) | $\sigma_j = \begin{cases} \frac{N+1}{d(2N-3)} & \text{if } j = 0 \\ \frac{2N}{d(2N-3)} & \text{if } 1 \leq j \leq d-3 \\ \frac{2N-1}{d(2N-3)} & \text{if } j = d-2 \\ \frac{3(N-d)}{d(2N-3)} & \text{if } j = d-1 \end{cases}$ |
| Moran process on a group splitting model | $\sigma_j = \begin{cases} \frac{n}{d(m+n-2)} & \text{if } 0 \leq j \leq d-2 \\ \frac{n+d(m-2)}{d(m+n-2)} & \text{if } j = d-1 \end{cases}$ |

### 3 Containment order

#### 3.1 A sufficient condition leading to the containment order

Consider two population structures *S*_1_ and *S*_2_ characterized by the (normalized) structure coefficients σ_1_ and σ_2_, and associated random variables *J*_1_ and *J*_2_, respectively. Let us also define the sequence of “gains from flipping”:

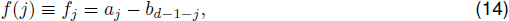

i.e., the gains in payoff experienced by a focal *B*-player interacting with *j A*-players (and *d — 1 — j B*-players) after all players in the group, including the focal, flip their strategies (*A*-players become *B*-players and vice versa).

Because of condition (2), the gains from flipping are increasing. Hence, a sufficient condition for *S*_1_ to be greater than *S*_2_ in the containment order is that

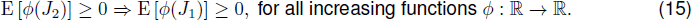

A sufficient condition for this is that

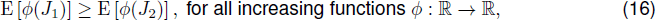

which is fulfilled by definition if *J*_1_ is greater than *J*_2_ in the (usual) stochastic order, denoted by 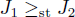 (ref. (8), p. 4).

There are many conditions leading to the stochastic ordering of two random variables (ref. (8), ch. 1). For instance, it is known that 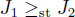 if and only if (ref. (8), p. 4)

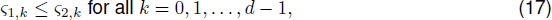

where ς is the distribution function corresponding to σ, i.e.,

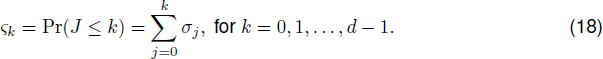

A simple sufficient condition leading to the set of inequalities given by (17) and hence to 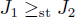 is that *S^-1^(σ_1_ – σ_2_)* (where *S*^-^(*a*) is the number of sign changes of the sequence *a*) and the sign sequence is –, + (ref. (8), p. 10). That is, if the structure coefficients σ_1_ “put more weight” in larger values of *j* than the structure coefficients σ_2_, then 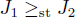. We summarize this observation in the following proposition.

**Proposition 1** (A sufficient condition leading to the containment order). *Let S_1_ and S_2_ be two population structures with (normalized) structure coefficients* σ_1_ and σ_2_, *respectively. If* S^-^(σ_1_ – σ_2_) = 1 *and the sign sequence is* –, +, *then* 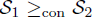.

#### 3.2 A sufficient condition leading to the incomparability in the containment order

Given two population structures *S*_1_ and *S*_2_, it could be that neither 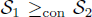 nor 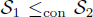 holds true. We are also interested in establishing a simple sufficient condition leading to such incomparability in the containment order, that we denote by 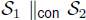. In order to derive this, suppose that the structure coefficients of *S*_1_ and *S*_2_ cross each other twice, i.e., that

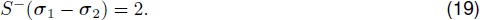

Condition (19) implies that *S^-^*(*ς*_1_ – *ς*_2_) (ref. (15), p. 621) and hence that (17) does not hold true. This in turn implies that *J*_1_ and *J*_2_ are incomparable in the stochastic order, i.e., *J*_1_ ||_st_ *J*_2_. Showing that (19) also implies 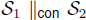 however requires some additional arguments. Indeed, note that *J*_1_ ||_st_ *J*_2_ does not necessarily imply 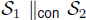: the stochastic order is a sufficient but not a necessary condition leading to the containment order (cf. (15) and (16)).

In order to prove that (19) leads to 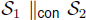, we make use of two other stochastic orders: the increasing convex order and the increasing concave order (ref. (8), p. 181). A random variable *J*_1_ is said to be greater than *J*_2_ in the increasing convex (resp. concave) order, denoted by 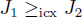 (resp. *J*_1_ ≥_icv_ *J*_2_), if

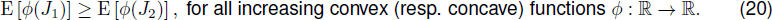

A simple condition leading to these orders is given in the following lemma.

**Lemma 1** (A sufficient condition leading to the increasing convex (resp. concave) order). *Let X and Y be two random variables with density functions p and q respectively. If*

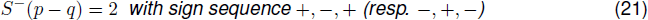

*then* 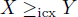 (*resp*. *X* ≥_icv_ *Y*).

*Proof*. Denote by *P* and *Q* the distribution functions associated to *X* and *Y*, respectively. Condition (21) implies (ref. (15), p. 621)

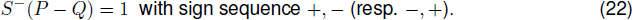

Since (22) implies the increasing convex (resp. concave) order (ref. (8), p. 194), this completes the proof.

Let us now consider two population structures *S*_1_ and *S*_2_ whose structure coefficients satisfy (19). Without loss of generality, suppose that the sign pattern is +,–, +. By Lemma 1, it follows that 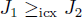 and 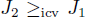. This suggests that there might be both (i) games with increasing and convex gains from flipping *f_j_* for which *S*_1_ (but not *S*_2_) fulfills the selection condition (5), and (ii) games with increasing and concave gains from flipping *f_j_* for which *S*_2_ (but not *S*_1_) fulfills the selection condition (5).

As an example of such games, consider a club goods game between cooperators (*A*) and defectors (*B*), where cooperators pay a cost *c > 0* in order to provide an excludable collective good that only cooperators can use, while defectors refrain from contributing and hence from using the good (7). This game is characterized by the payoff sequences:

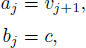

where *v_k_* gives the value of the collective good as a function of the total number of cooperators, *k = j + 1*, and *c* is the payoff defectors obtain. We further assume that *v_k_* is given by

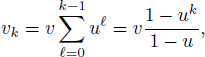

where *v > 0* is some baseline value, and *u > 0* is a synergy or discounting parameter (3). Furthermore, we require that *v/c > γ*, where 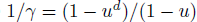, so that (3) is fulfilled.

The gains from flipping of this game are then given by

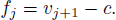

Let us first impose the condition:

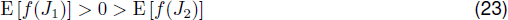

so that *S*_1_ but not *S*_2_ satisfies the selection condition (5). Condition (23) is satisfied if

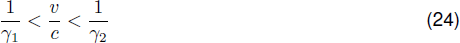

where

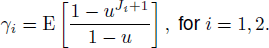

Note that (24) is satisfied if *u >* 1, because in this case 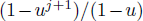 is increasing and convex and 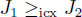. Additionally, 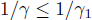 always holds.

Let us now impose the condition:

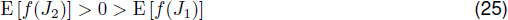

so that *S*_2_ but not *S*_1_ satisfies the selection condition (5). In this case, (25) is satisfied if

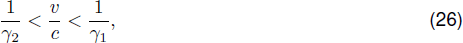

which holds true if *0 < u < 1*, as in this case 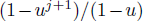) is increasing and concave and 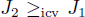. Additionally, 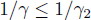 always holds.

We summarize the previous observations in the following proposition.

**Proposition 2** (A sufficient condition leading to the incomparability in the containment order). *Let S*_1_ *and S*_2_ *be two population structures with (normalized) structure coefficients* σ_1_ *and* σ_2_, *respectively*. *If S*^−^ (σ_1_ – σ_2_) = 2 *then* 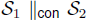, *i.e., neither* 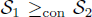 *nor* 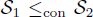 *hold true. Moreover, if the sign sequence of* σ_1_ – σ_2_ *is* +, –, + (*resp*. –, +, –*) then it is possible to find cooperation games with convex (resp. concave) gains from flipping *f_j_* such that the selection condition* (5) *is satisfied for S*_1_ *but not for S*_2_ (*resp. for S*_2_ *but not for S*_1_) *and cooperation games with concave (resp. convex) gains from flipping f_j_ such that the selection condition* (5) *is satisfied for S*_2_ *but not for S*_1_ (*resp. for S*_1_ *but not for S*_2_).

#### 3.3 The containment order is a total order for *d = 2* but a partial order for *d ≥ 2*

Propositions 1 and 2 allow us to prove the following result.

**Proposition 3** (The containment order is total for *d = 2* but partial for *d = 3*). *Consider the set of all possible population structures* {*S*} *for a given group size d*. {*S*} *is totally ordered under* ≤_con_ *for d* = 2 *but only partially ordered under* ≤_con_ *for d ≥ 3*.

*Proof*. For *d* = 2, the probability mass function given by the normalized structure coefficients σ consists of only two points. Consequently, σ_1_ – σ_2_ has either (i) no sign changes (i.e., σ_1_ = σ_2_), which implies *S*_1_ =_con_ *S*_2_; (ii) a sign change from – to +, which implies 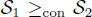; or (iii) a sign change from + to –, which implies 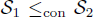. For *d* ≥ 3, the probability mass function given by the normalized structure coefficients σ consists of *d* ≥ 3 points. In this case, it is always possible to find *S*_1_ and *S*_2_ such that σ_1_ – σ_2_ has two sign changes. In this case, neither 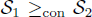 nor 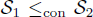 hold true.

#### 3.4 Examples

In the following, we state several results concerning the containment order for the population structures listed in Table 1. We omit the proofs, as they are straightforward applications of Propositions 1 and 2 above.

**Proposition 4** (Containment order for well-mixed populations updated with a Moran process). *Denote by* 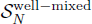 *a well-mixed population of size N with a Moran process as updating rule. Then* 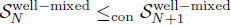 *for all N ≥ d*.

**Proposition 5** (Containment order for cycles updated with a Moran death-Birth process). *Denote by* 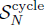 a cycle of size N with death-Birth updating. Then 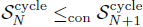 for all N > d.

**Proposition 6** (Containment order for cycles and well-mixed populations updated with a Moran death-Birth process). Let 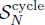 and 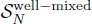 be respectively a cycle and a well-mixed population of size N, both updated with a Moran death-Birth process. Then, for all d and all N > d, 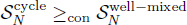.

**Proposition 7** (Containment order for group splitting models and well-mixed populations, both updated with a Moran death-Birth process). *Let* 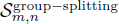 *be a group splitting model with m groups of maximum size n and rare group splitting (q ≪ 1), and* 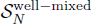 *a well-mixed population of size N, both updated with a Moran death-Birth process. We have that* 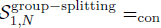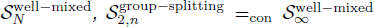 *for any n, and* 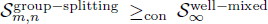 *for m ≥ 3 and any n*.

**Proposition 8** (Containment order for cycles and group splitting models, both updated with a Moran death-Birth process). *Let* 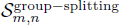 *be a group splitting model with m groups of maximum size n and rare group splitting (q ≪ 1), and S^cycle^ be a cycle of size N, both updated with a Moran death-Birth process. In the limit of large N = mn we have:*

1. 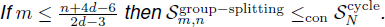
2. 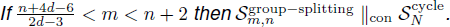
3. 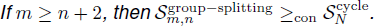

**Proposition 9** (Aspiration dynamics vs. Moran process in well-mixed populations). *Let S^aspiration^ and* 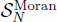 *be well-mixed populations of size N ≥ d, updated with aspiration dynamics and a Moran process, respectively. We have:*

1. 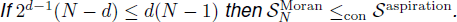
2. 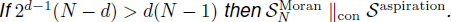

### Volume order

In the following, we give a formula for the volume of cooperation games, as defined by inequalities (2)-(4). For this, we find convenient to define 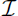 as the polytope given by inequalities (2) and (4), and 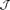 as the polytope given by inequalities (2), (4), and 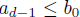 (which is the opposite of (3)). The volumes of these two polytopes are easy to calculate exactly using probabilistic arguments. Indeed, we have the following two lemmas.

**Lemma 2** (Volume of 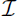). *We have that*

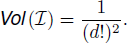

*Proof*. Calculating the volume of 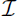 is equivalent to calculating the probability that two sequences 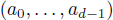 and 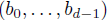, with elements randomly and independently drawn from the interval [0, 1], are such that 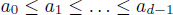 and 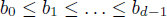. For each sequence, the probability of having a randomly ordered sequence of length *d* is 1/*d*!, since *d*! is the number of permutations of *d* distinct objects and only one of such permutations will be given in the specified order. Since the two sequences are independent, the total probability is given by 1/(*d*!)^2^.

**Lemma 3** (Volume of 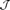). *We have that*

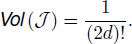

*Proof*. Calculating the volume of 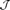 is equivalent to calculating the probability that the sequence

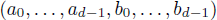

with elements randomly and independently drawn from the interval [0, 1] is such that *a*_0_ ≤ *a*_1_ ≤ … ≤ *a*_*d*–1_ ≤ *b*_0_ ≤ *b*_1_ ≤ … ≤ *b*_*d*–1_. Following the same argument as in the proof of Lemma 2, this probability is equal to 1/(2*d*)!.

Making use of these two lemmas, we can find an expression for the volume of *P* the polytope of cooperation games. We state this result in the following proposition.

**Proposition 10** (Volume of cooperation games). *The volume of cooperation games is given by*

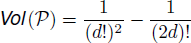

*Proof*. Follows from Lemmas 2 and 3 upon noticing that *P* = 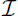 – 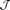

Some population structures, such as large well-mixed populations updated with a Moran process, and finite well-mixed populations updated with the aspiration dynamics, are such that their structure coefficients are symmetric, i.e., *σ_ό_* = *σ*_*d*-1-*j*_ for all *j*. We are intereted in calculating the fraction of cooperation games for which such population structures favor strategy *A*. In order to calculate this result, we need the following lemma. The lemma may appear to be obvious for symmetry reasons, but the additional requirement that we are dealing with cooperation games (and hence a subset of the hypercube of all possible games) adds a further complication.

**Lemma 4.** *Let S be a population structure with symmetric structure coefficients, i.e., σ*_*j*_ = *σ*_*d*–1–*j*_ *for all j. Let also* 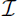_+_ *be the subset of* 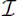 *for which A is favored over B. Then*

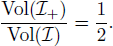

*Proof*. Denote by 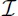_–_ and 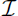_0_ the subsets of 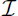 such that 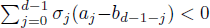 and 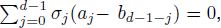 respectively. Then we arrive at a partition of set 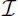, namely 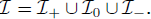 Since there must exist a *j*\* such that *σ_j*_* ≠ 0, the solution space of 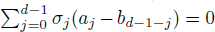 is of lower dimension than 2*d*, and hence Vol(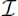_0_) = 0. Thus Vol(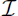) = Vol(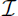_+_) + Vol(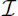I_–_).

In the following, we prove that Vol(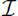_+_) = Vol(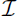_–_). For this, we define the mapping

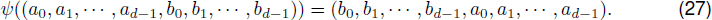

For every 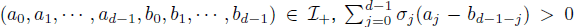 holds, which implies that 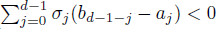 holds. Since *σ_j_* = *σ*_*d*–1–*j*_, we have 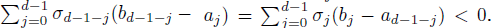 This implies that 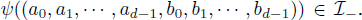 Thus 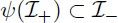. Similarly, we have 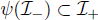. Therefore 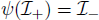. This leads to

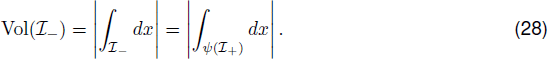

In addition, since *ψ*^2^ is the identity mapping (*ψ*^2^ = *I*), *ψ* is invertible and the inverse mapping is the mapping itself, i.e., *ψ*^−1^ = *ψ*. This leads to

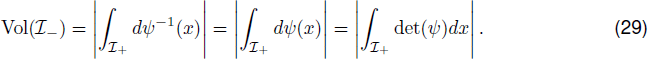

Here det(*ψ*) is the determinant of the Jacobian matrix of the transformation *ψ* at *x*. Further, considering that *ψ* is a linear mapping, *ψ*^2^ = *I* implies | det(*ψ*)| = 1. Thus (29) yields

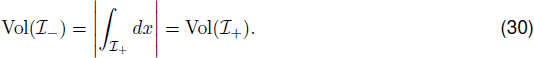

Therefore Vol(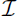) = Vol(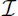_+_) + Vol(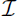_–_) = 2Vol(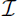_+_), or

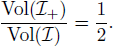

With this lemma, we can prove the following proposition.

**Proposition 11.** *Let S be a population structure with positive symmetric structure coefficients, i.e., σ_j_ = σ_d–1–j_ for all j, and Q the polytope asociated to all cooperation games for which A is favored over B under S. Then*

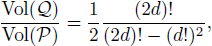

which is a decreasing function of *d*, and is equal to 1/2 in the limit of large *d*.

*Proof*. It is easy to check that for every 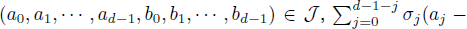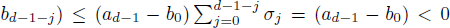 holds true, which implies 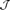 ⊂ 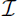_–_. Since 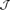 ⊂ 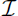_–_ and 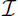 = 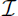_–_ ∪ 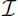_0_ ∪_+_, then *Q* = 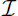_+_. Moreover *P* = 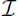 – 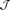. Hence

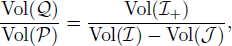

and by Lemma 4

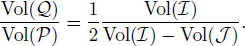

Using Lemmas 2 and 3, we finish the proof.

